# In silico analysis of predicted differential MHC binding and CD8+ T-cell immune escape of SARS-CoV-2 B.1.1.529 variant mutant epitopes

**DOI:** 10.1101/2022.01.31.478157

**Authors:** Pablo Riesgo-Ferreiro, Ranganath Gudimella, Thomas Bukur, Patrick Sorn, Thomas Rösler, Barbara Schrörs, Martin Löwer

## Abstract

**Introduction:** The B.1.1.529 (Omicron) SARS-CoV-2 variant has raised global concerns due to its high number of mutations and its rapid spread. It is of major importance to understand the impact of this variant on the acquired and induced immunity. Several preliminary studies have reported the impact of antibody binding and to this date, there are few studies on Omicron’s CD8^+^ T-cell immune escape.

**Methods:** We first assessed the impact of Omicron and B.1.617.2 (Delta) variant mutations on the SARS-CoV-2 spike epitopes submitted to the Immune Epitope Database (IEDB) with positive outcome on MHC ligand or T-cell assays (n=411). From those epitopes modified by a mutation, we found the corresponding homologous epitopes in Omicron and Delta. We then ran the netMHCpan computational MHC binding prediction on the pairs of IEDB epitopes and matching homologous epitopes over top 5 MHC I alleles on some selected populations. Lastly, we applied a Fisher test to find mutations enriched for homologous epitopes with decreased predicted binding affinity.

**Results:** We found 31 and 78 IEDB epitopes modified by Delta and Omicron mutations, respectively. The IEDB spike protein epitopes redundantly cover the protein sequence. The WT pMHC with a strong predicted binding tend to have homologous mutated pMHC with decreased binding. A similar trend is observed in Delta over all HLA genes, while in Omicron only for HLA-B and HLA-C. Finally, we obtained one and seven mutations enriched for homologous mutated pMHC with decreased MHC binding affinity in Delta and Omicron, respectively. Three of the Omicron mutations, VYY143-145del, K417N and Y505H, are replacing an aromatic or large amino acid, which are reported to be enriched in immunogenic epitopes. K417N is common with Beta variants, while Y505H and VYY143-145del are novel Omicron mutations.

**Conclusion:** In summary, pMHC with Delta and Omicron mutations show decreased MHC binding affinity, which results in a trend specific to SARS-CoV-2 variants. Such epitopes may decrease overall presentation on different HLA alleles suggesting evasion from CD8+ T-cell responses in specific HLA alleles. However, our results show B.1.1.529 (Omicron) will not totally evade the immune system through a CD8^+^ immune escape mechanism. Yet, we identified mutations in B.1.1.529 (Omicron) introducing amino acids associated with increased immunogenicity.

**Availability:** All the code and results from this study are available at https://github.com/TRON-bioinformatics/omicron-analysis.

## Introduction

The first sequences of the SARS-CoV-2 B.1.1.529 lineage were uploaded to GISAID [1] on 23^rd^ November 2021 from Botswana and South Africa, although its origin is still unclear. It raised a lot of attention due to its high number of mutations in the spike protein, and hence, it has potential impact on the naturally acquired and induced immunity. The World Health Organization classified B.1.1.529 as a Variant of Concern (VOC) on 26 November 2021 and named it as Omicron [2].

Scientists raced to investigate two lines of work: 1) viral transmission and infection rate 2) the effectivity of the SARS-CoV-2 vaccines. Preliminary evidence on virulence and increase in rate of infection came from South Africa where the variant was first reported [3]. There are reports of increased binding to the human ACE2 protein (14 results in BioRxiv for the term ‘SARS-CoV-2 omicron ACE2’ on 14 December). Recent reports suggest structural changes in new spike protein receptor binding domain (RBD), show reduced antibody interaction [4] and demonstrate Omicron to be escaping RBD neutralizing antibody drugs such as LY-CoV016/LY-CoV555 cocktail, REGN-CoV2 cocktail, AZD1061/AZD8895 cocktail, and BRII-196 [5]. Preliminary results published by Pfizer and BioNTech show that vaccine neutralization was reduced on Omicron and a third dose of the vaccine was recommended [6].

While B-cells form a first line of defense, CD8^+^ T-cells are a second line of defense that identify and kill infected host cells. High levels of CD8^+^ T-cells have been associated to a less severe disease [7]. A recent preprint found little overlap of the CD8^+^ T-cell epitopes (n=52) from patients (n=30) recovered from the Beta strain with the Omicron mutations [8] and a recent publication confirms that a majority of known SARS-CoV-2 epitopes are not affected by mutations and will still support CD8+ T-cell activity in omicron [9]. This confirmed from recent studies on vaccinated and omicron infected patients which report slight reduction (15% to 20%) of CD8+ T cell response and SARS-CoV2 vaccines are extensively cross-reactive with Omicron variant despite of large number of mutations [10–12]. In this study, we aim at understanding how the new spike protein mutations in B.1.1.529 may be a CD8+ T-cell immune escape mechanism via a decreased epitope MHC binding affinity.

The IEDB database [13] holds a list of 411 SARS-CoV-2 spike protein epitopes with a positive outcome to T-cell and MHC ligand assays. These epitopes obtained from the original SARS-CoV-2 Alpha lineage or from some of the strains that became dominant at any geography and time. Following the same approach as in our previous SARS-CoV-2 study [14] we use this list from IEDB as our truth set.

The 34 mutations in the Omicron spike protein hit 78 IEDB epitopes, from which we obtained the corresponding homologous mutated sequences. We selected the five most frequnet HLA-A, HLA-B and HLA-C alleles for selected host populations from the Allele Frequency Net Database [15]. Using netMHCpan 4.1 [16], we predicted MHC binding affinity for the pairs of IEDB epitopes – mutated epitopes for each of the alleles. Finally, we tested each mutation for enrichment of epitopes with a decreased predicted binding affinity with the aim of pinpointing those mutations most involved in CD8^+^ immune escape.

We found that for those pMHC with strong predicted MHC binding, the majority of corresponding B.1.1.529 mutated pMHC showed a decreased MHC binding. On the other hand, we did not detect a strong signal of decreased binding in B.1.1.529 for HLA-A, although it does in B.1.627.2. A similar trend was observed in all studied populations. Finally, we identified seven mutations enriched for epitopes with decreased binding. Aromatic and large amino acids have been associated to immunogenicity before [17], B.1.1.529 has three mutations replacing such amino acids (VYY143-145del, K417N and Y505H) which are significantly enriched for decreased MHC binding affinity.

## Results

We aim at understanding how the mutations in B.1.1.529, designated as Omicron by the WHO, may be involved in CD8^+^ T-cell immune escape mechanisms through a decreased MHC binding affinity of epitopes derived from the spike protein. We take as a baseline the 411 SARS-CoV-2 epitopes with a positive outcome in MHC ligands or T-cell assays restricted to MHC I from IEDB database [13] (queried on 5^th^ January 2022). We identify those IEDB epitopes modified by mutations in B.1.1.529 and identify its homologous counterparts. Further, we analyze the differential predicted MHC binding rank according to netMHCpan 4.1 [16]. Lastly, we identify those mutations enriched for mutated pMHC with a decreased MHC binding.

### B.1.627.2 and B.1.1.529 spike protein mutations and IEDB epitopes

The Omicron mutations were listed in the pango-designation repository on 23 November 2021 (https://github.com/cov-lineages/pango-designation/issues/343). The spike protein has 34 mutations; this represents a quantitative increase from the nine mutations in the current prevalent Delta strain B.1.627.2. Some of these mutations modify the IEDB epitopes of interest. In Figure *1*, we observe that B.1.627.2 and B.1.1.529 share with the reference respectively 367 (92.21 %) and 320 (80.40 %) IEDB epitopes. There are 13 epitopes without an exact match in the reference genome; of these, the one epitope present only in the Omicron sequence overlaps the mutation K417N that is part of the Beta lineage too. The 31 and 78 IEDB epitopes from the Wuhan strain and mutated in either B.1.627.2 or B.1.1.529, respectively, are of special interest.

**Figure 1:**
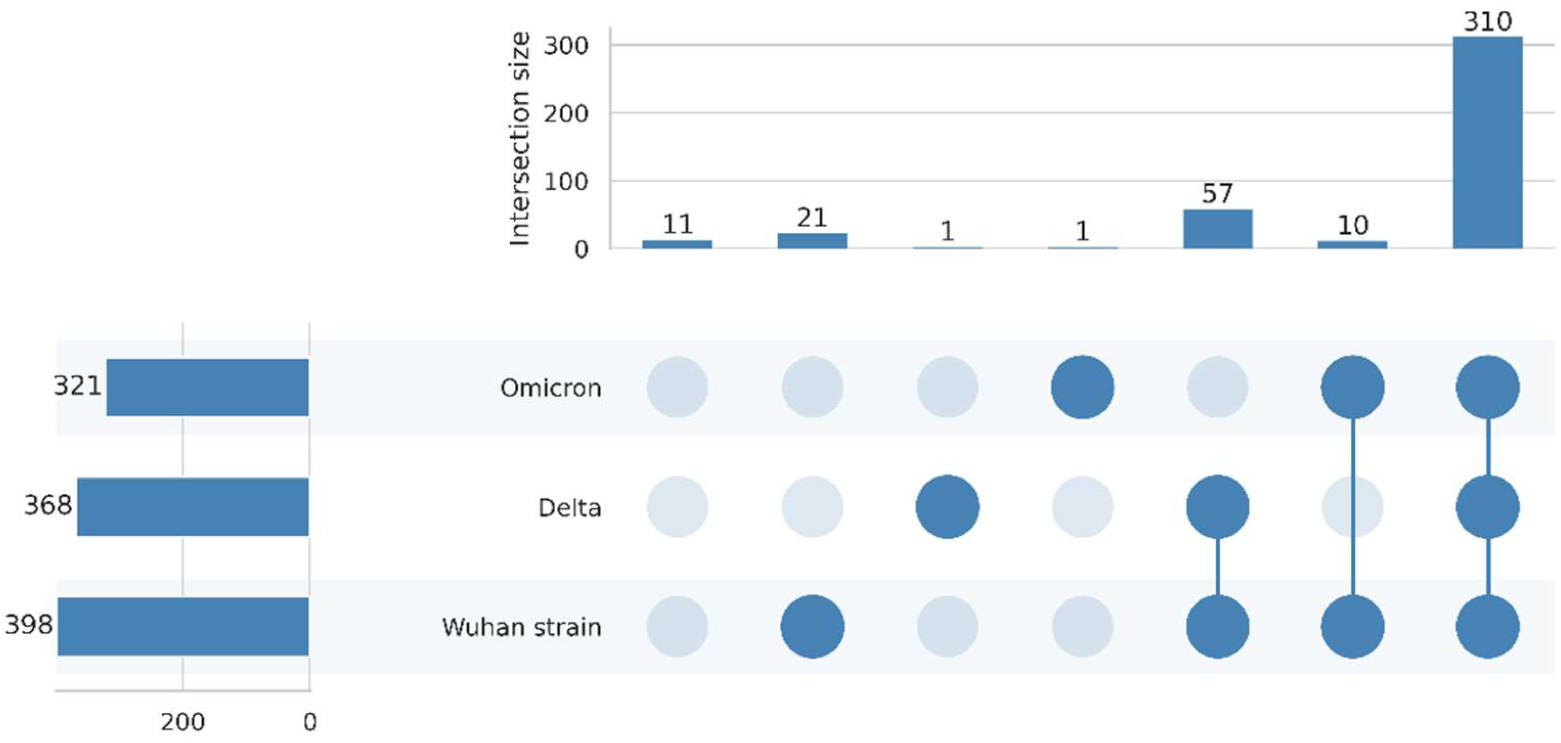
IEDB epitopes from SARS-CoV-2 spike glycoprotein with a positive outcome to T-cell and MHC ligand assays restricted to MHC I presence in Omicron, Delta and the Wuhan strains. The majority of the IEDB epitopes are present in the three strains. Sequences having an exact match found by performing a global alignment as described in the methods section. Data downloaded from IEDB.org on January 5, 2022.

The IEDB epitopes overlap the SARS-CoV-2 spike protein redundantly; i.e. more than one epitope may overlap a given position. In addition, as shown in Figure *2* the epitope coverage is not homogenous through the spike protein. L452R is the B.1.627.2 mutation mutating most epitopes, nine; while T19R, T478K and D950N only mutate one epitope (see supplementary table 1). In the case of B.1.1.529 S371L hits the most epitopes, nine; and T547K and N969K do not mutate any epitope. There are three shared mutations between B.1.627.2 and B.1.1.529: G142D, T478K and D614G. Last, there is a mutation hitting the same amino acid in both strains: P681R in B.1.627.2 and P681H in B.1.1.529. The relation between the number of mutations and the total number of mutated epitopes is a monotonically increasing non-linear relationship.

**Figure 2:**
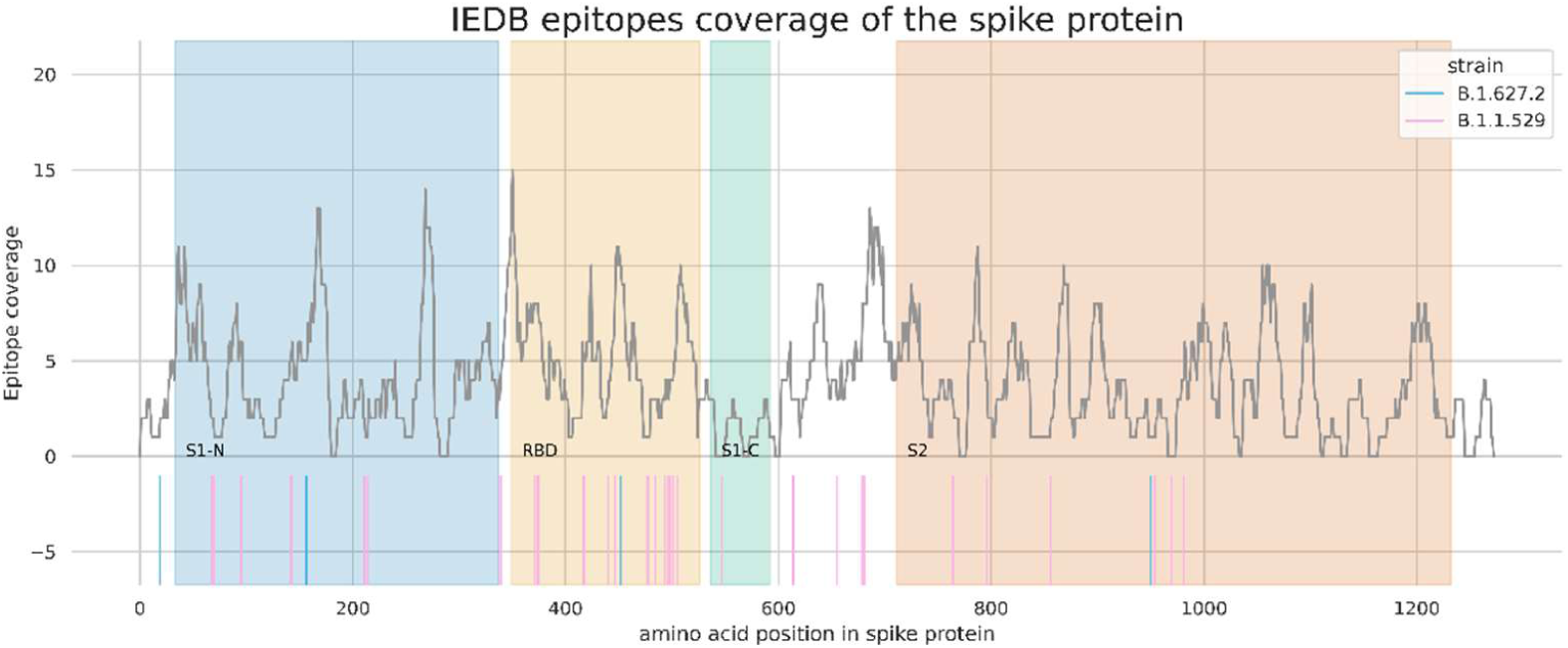
Number of IEDB epitopes overlapping each position of the spike protein (grey line) and B.1.627.2 and B.1.1.529 mutation positions (blue and pink rug plots). There are 12 gaps not covered by any epitope within the protein with a minimum and maximum length of three and 10 amino acids. Two, two and six gaps lie within the subunit 1 N-terminal domain, subunit 2 C-terminal domain and subunit 2, respectively. The RBD is fully covered by IEBD epitopes.

### Decreased predicted MHC binding observed in B.1.1.529

Those IEDB epitopes hit by a mutation have a corresponding homologous epitope in B.1.1.529. We compared the netMHCpan predicted MHC binding affinity from these pairs of WT and mutated epitopes in both B.1.627.2 and B.1.1.529 on the top 5 most frequent alleles of the classic MHC I genes from different populations (other populations available in supplementary figures 1, 2, 3 and 4). We observe in Figure *3* that the triplets (consisting of IEDB epitope, homologous epitope and HLA allele) with a decreased predicted MHC binding affinity (quotient < 0.8) are enriched in the subset of IEDB epitopes with a strong WT MHC binding prediction (netMHCpan rank < 10). Enrichment of overall decreased binding in the IEDB pMHC with increased binding show Fisher test p-values of B.1.1.529 are 3.24×10^−8^, 9.1×10^−5^ and 1.19×10^−9^ for Sub-Saharan, black South African and European populations. While Fisher test, p-values of B.1.627.2 show less significance i.e. 1.03×10^−2^, 9.71×10^−4^ and 9.33×10^−3^ for Sub-Saharan, black South African and European populations.

Next, we focus on those pMHC with a strong predicted MHC binding. We set two different thresholds to consider a MHC binding affinity strong and very strong, a rank of 10 and a rank of one, respectively. Then we explored the distribution of the quotient of predicted binding affinity between WT and B.1.1.529 epitopes across different chosen populations (other population results in supplementary figures 5 and 6). We observed for HLA genes B and C a decreased predicted MHC binding affinity, which is more marked for the more conservative threshold (Figure *4*). On the other hand, gene A shows a more homogenous distribution indicating neither increased or decreased predicted binding.

**Figure 3:**
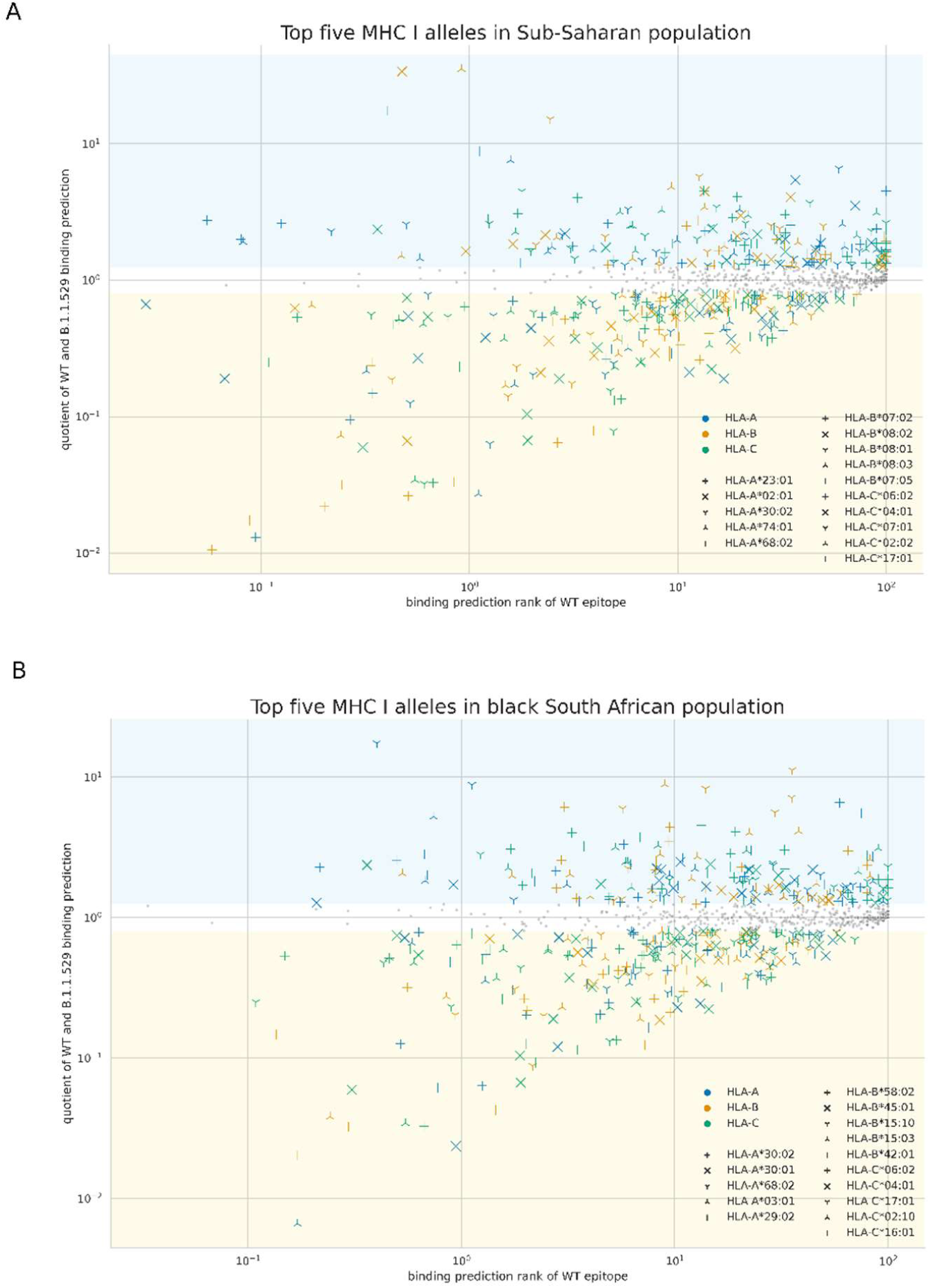
Differential MHC binding of the IEDB epitope-homologous epitope pairs in B.1.1.529. On the X-axis the predicted MHC binding rank in the IEDB epitope and on the Y-axis the quotient between the predicted MHC binding rank in the IEDB epitope and the B.1.1.529 homologous epitope. The yellow shaded area indicates decreased predicted binding (quotient < 0.8) and the blue shaded area indicates increased binding (quotient > 1.25). A: top five MHC I alleles in Sub-Saharan population; B: top five MHC I alleles in black South African population.

**Figure 4:**
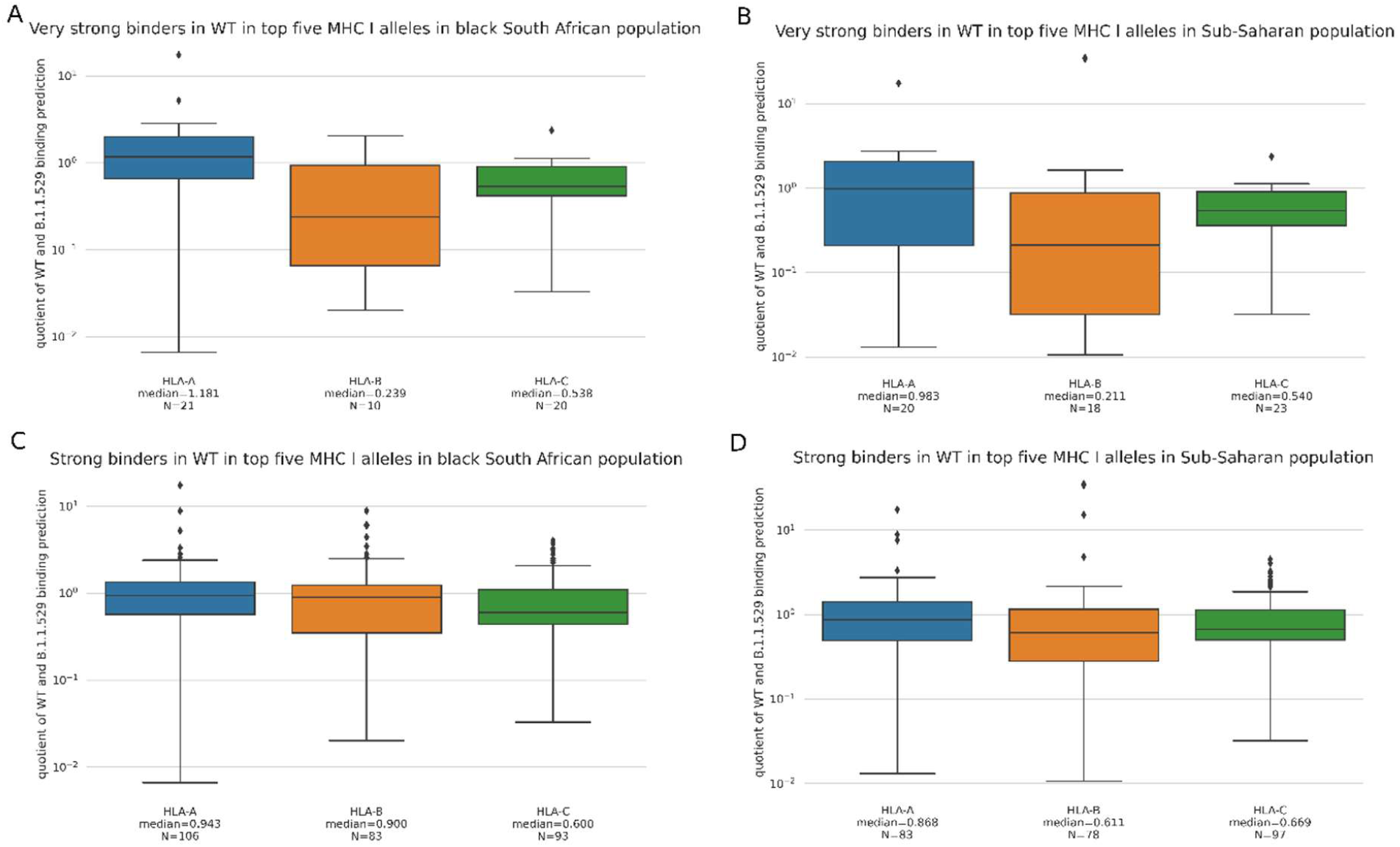
Distribution of quotient of predicted MHC binding in the WT and B.1.1.529. A: epitopes with WT binding rank < 1.0 on top five MHC I alleles in black South African population; B: epitopes with WT binding rank < 1.0 on top five MHC I alleles in Sub-Saharan population; C: epitopes with WT binding rank < 10.0 on top five MHC I alleles in black South African population; D: epitopes with WT binding rank < 10.0 on top five MHC I alleles in Sub-Saharan population.

### Comparison to B.1.627.2

The comparison between B.1.627.2 and B.1.1.529 is important as B.1.627.2 is the current prevalent strain and not the reference genome isolated at the end of 2019. There is a quantitative difference in the number of mutations and hence of modified IEDB epitopes. To understand if the IEDB epitopes modified by B.1.1.529 and B.1.627.2 mutations show a different pattern we looked at the distribution of quotients (*Figure 4 & Figure 5*). We confirmed the expected quantitative difference (21, 10 and 20 IEDB epitopes modified by B.1.1.529 mutations with a predicted MHC binding rank below 1.0 in the Sub-Saharan population as opposed to 10, 4 and 3 modified by B.1.627.2). Although there are fewer epitopes in B.1.627.2, the HLA A gene epitopes show a decreased predicted MHC binding affinity with a median rank of 0.439, which is not observed in B.1.1.529. HLA B and C show similar values to B.1.1.529. This trend was found in all populations under scrutiny.

**Figure 5:**
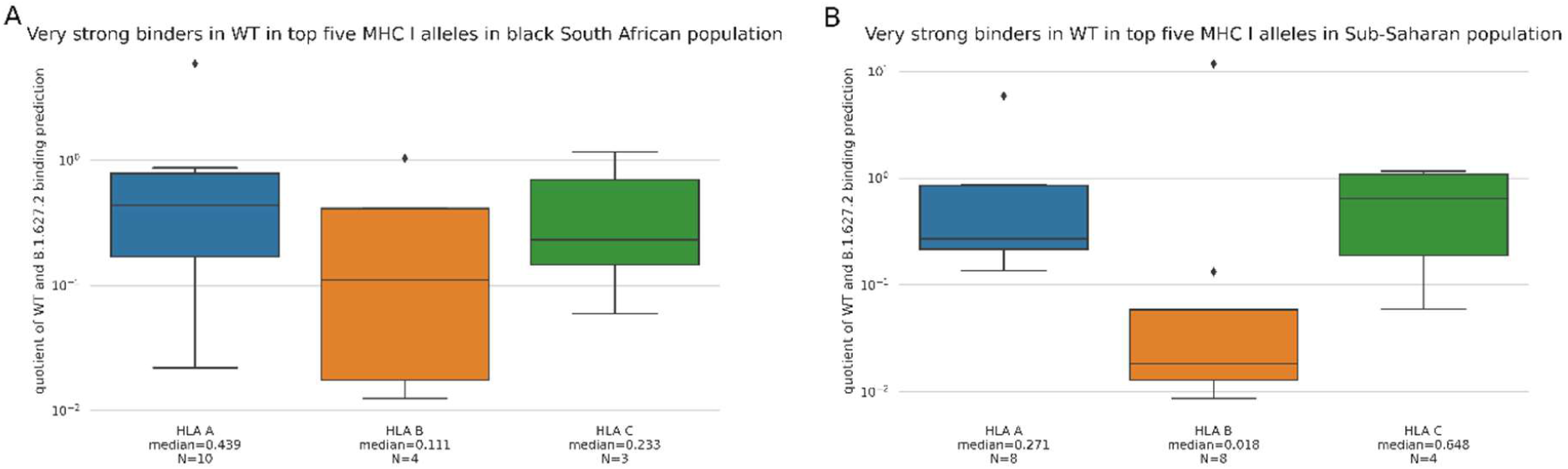
Distribution of quotient of predicted MHC binding in the WT and B.1.627.2. A: top 5 HLA alleles in the Sub-Saharan population for WT binding rank < 1.0; B: top 5 HLA alleles in the black South African population for WT binding rank < 1.0

Furthermore, when we tested for enrichment of decreased predicted MHC binding affinity (quotient < 0.8) in the subset of IEDB epitopes with a strong WT MHC binding prediction (netMHCpan rank < 10) in B.1.627.2 we found less significant values, Fisher p-values of 0.01, 0.001 and 0.009 for Sub-Saharan, black South African and European populations respectively.

### Prediction of CD8^+^ T-cell immune escape mutations

Predicting whether a SARS-CoV-2 strain has better CD8^+^ T-cell immune escape than previous strains is challenging when assessing the epitopes. The signal may be distributed across multiple epitopes, HLA genes and alleles; and there are other immune escape mechanisms than HLA binding affinity of epitopes. Furthermore, a given homologous epitope may be overlapped by more than one mutation. We tested every mutation in the spike protein of each lineage for enrichment of mutated pMHC with a decreased predicted binding (see methods). We defined decreased predicted binding as those WT epitopes with strong predicted MHC binding affinity (rank below 10) and quotient between predicted binding of WT and mutated epitopes below 0.8. We did this across the five most frequent MHC I alleles of several populations (as defined in http://www.allelefrequencies.net): Sub-Saharan, black South African, white South African, Zulu South African and European. After removing repeated alleles between populations this lead to 35 HLA alleles across HLA-A, HLA-B and HLA-C. Considering mutations with a p-value below 0.0001 and an odds ratio above 1.0 we found one single amino acid substitution (D614G) enriched in B.1.627.2 (Figure *6* A). This mutation lies between the S1 and S2 subunits. Additionally, in B.1.1.529 we found two deletions and one insertion in the N-terminal domain (VYY143-145del, 214EPE and N211+L212I) and three single amino acid substitutions (SNVs) (Figure *6* B). Two of the SNVs (K417N and Y505H) lie in the RBD. D614G is a shared mutation between both lineages and it is also enriched in B.1.1.529.

**Figure 6:**
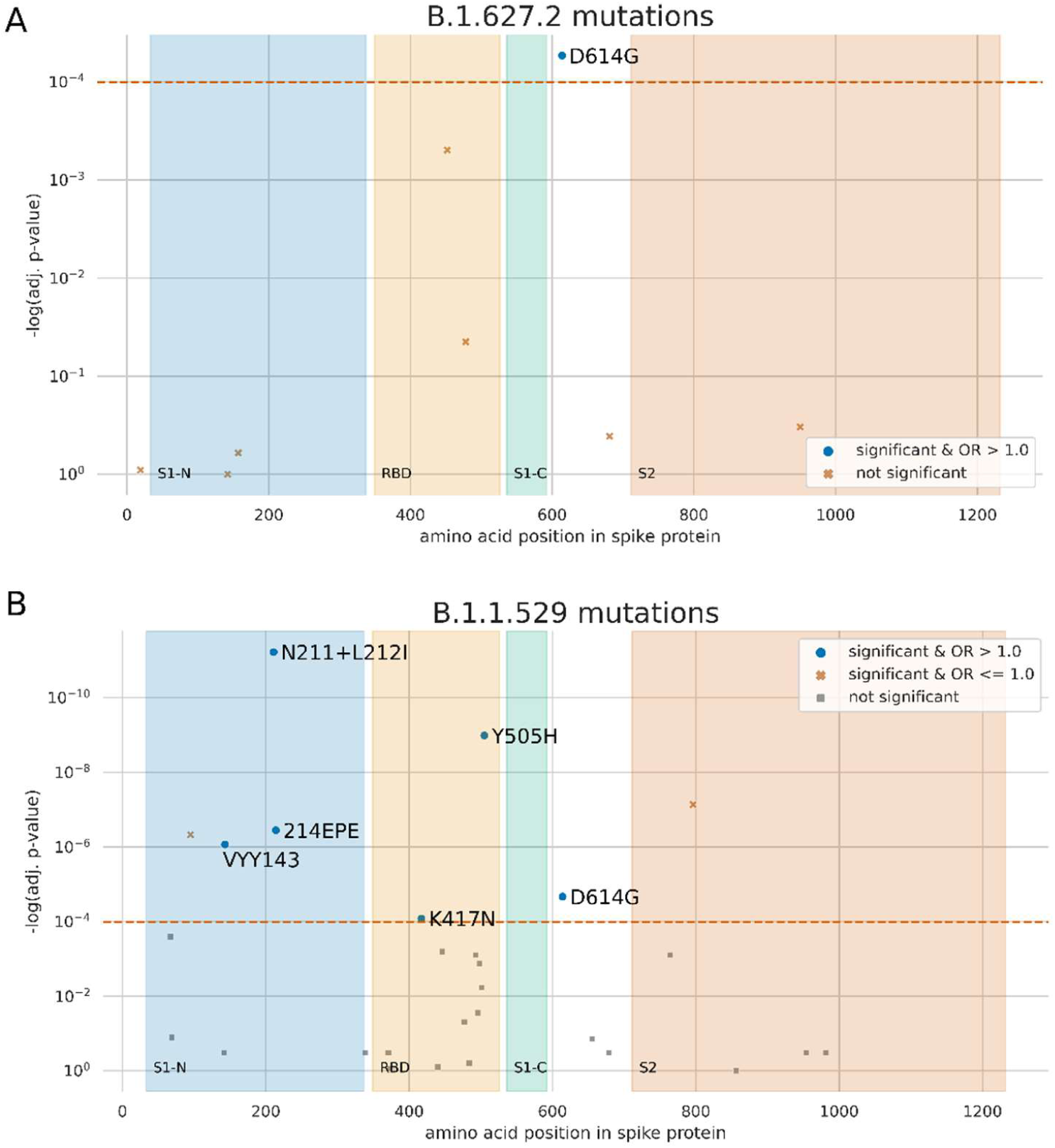
Enrichment analysis of mutations on the subset of strong binders in WT with decreased predicted MHC binding for the homologous epitopes. The X-axis is the position in the spike protein. The different spike domains are shaded in blue, orange, green and red. The Y-axis has the median Fisher exact test p-value log transformed and reversed. Enriched mutations with an odds ratio greater than 1.0 are labelled. For visualization purposes, a pair of mutations in close by amino acids with similar enrichment results are merged. A: B.1.627.2; B: B.1.1.529

Two mutations in B.1.1.529 remove aromatic Tyrosine (Y) amino acids, VYY143 and Y505H, both of these mutations are significantly enriched in mutated pMHC with a decreased binding (adjusted p-values: 8.65×10^−7^ and 1.03×10^−9^). Y505H is not present in any of the dominant lineages, although according to GISAID data [1] processed through CoVigator [19] (manuscript in preparation) it has been observed before on 105 isolates; of which 60 in Ireland, 15 in the United Kingdom, five in France and five in Germany. K417N is also enriched and it replaces a Lysine (K) amino acid (adjusted p-value: 8.29×10^−5^), it has been observed before in the B.1.351 strain (ie: Beta). Conversely, there are two other mutations with a significant enrichment with odds ratio below 1.0 –enriched for non-decreased binding– T95I and D796Y introducing a large Isoleucine and a Tyrosine, respectively. Large and aromatic amino acids, like Lysine, Isoleucine and Tyrosine, have been associated before with increased immunogenicity [17].

## Methods

We downloaded the SARS-CoV-2 reference genome from ftp://ftp.ensemblgenomes.org/pub/viruses/fasta/sars_cov_2/cdna/Sars_cov_2.ASM985889v3.cdna.all.fa.gz. We downloaded the Pfam domains from the Ensembl annotations at ftp://ftp.ensemblgenomes.org/pub/viruses/json/sars_cov_2/sars_cov_2.json. We obtained the mutations in B.1.1.529 lineage from the following issue in pango designation GitHub repository https://github.com/cov-lineages/pango-designation/issues/343: A67V, Δ69-70, T95I, G142D/Δ143-145, Δ211/L212I, ins214EPE, G339D, S371L, S373P, S375F, K417N, N440K, G446S, S477N, T478K, E484A, Q493R, G496S, Q498R, N501Y, Y505H, T547K, D614G, H655Y, N679K, P681H, N764K, D796Y, N856K, Q954H, N969K, L981F. We obtained the mutations in the B.1.627.2 from the pango constellations site (https://github.com/cov-line-ages/constellations/blob/main/constellations/definitions/cB.1.617.2.json): T19R, G142D, L452R, T478K, P681R, D950N. We needed to add in the mutation D614G shared by alpha, beta, gamma, delta and omicron. Also, the deletion 22028:GAGTTCA>G was missing, this is reported in Wikipedia entry for B.1.627.2 lineage (https://en.wikipedia.org/wiki/SARS-CoV-2_Delta_variant) and we confirmed its existence in our CoVigator site that gathers information from ENA and GISAID (www.covigator.tron-mainz.de, manuscript in preparation). This deletion translates at the amino acid level in two mutations E156G and Δ157-158. We downloaded the SARS-CoV-2 IEDB MHC I epitopes with a positive outcome to T-cell response or MHC binding from the IEDB site http://www.iedb.org/ [13] on the 26^th^ November 2021.

We combined the spike protein reference sequence and the mutations from B.1.1.529 and B.1.627.2 into the alternative spike proteins of each lineage with custom code. We obtained the homologous epitopes by performing a global alignment using BioPython [18] with parameters mode=‘global’, match=2, mismatch=-2, open_gap_score=-3 and extend_gap_score=-1.

We selected the top five HLA alleles from some selected populations from the resource http://www.allelefrequencies.net. We fetch data from the following populations: Europe, Sub-Saharan Africa, Black in South Africa (n=142) (http://www.allelefrequen-cies.net/pop6001c.asp?pop_name=South%20African%20%20Black), white in South Africa (n=102) (http://www.allelefrequencies.net/pop6001c.asp?pop_name=South%20Af-rica%20Caucasians), Zulu in South Africa (n=100, only HLA-A and HLA-B available) (http://www.allelefrequencies.net/pop6001c.asp?pop_name=South%20Africa%20Natal%20Zulu).

MHC binding is performed with the tool netMHCpan 4.1 [16] with the following parameters ‘netMHCpan -p {fasta} -a {hla_allele} -s’ for every epitope of 8, 9, 10 and 11 amino acids using NeoFox [20] API. We keep only the rank of the best binding for every allele independently. We calculate the amplitude as the quotient between the IEDB epitope best rank and the homologous epitope best rank from either B.1.1.529 or B.1.627.2 as described in [21].

To test for enrichment of overall decreased binding in the IEDB pMHC with strong binding we apply the Fisher’s exact test on each population and lineage separately as indicated in the contingency table below (Table *1*). We did not adjust for multiple testing in this case as we only performed six tests.

**Table 1:** contingency table for Fisher’s test for the enrichment of overall decreased binding. There is one test per population for each Omicron and Delta lineages.

|  | Decreased binding<br>(quotient < 0.8) | Increased binding<br>(quotient > 1.25) |
| --- | --- | --- |
| Binding WT (rank < 10) | a | b |
| Not binding WT | c | d |

Finally, we test each mutation for the enrichment of mutated pMHC with decreased binding with the Fisher’s exact test as implemented in Scipy [22] and the contingency table indicated below (Table *2*). Even though the number of tests is limited to the number of mutations, we apply FDR Benjamini-Hochberg multiple test correction. We evaluate each mutation on all top five alleles of the populations described above without repeated alleles. Mutations with an odds ratio below 1.0 and a significant p-value correspond to those mutations with less decreased binding epitopes than expected by chance, while mutations with an odds ratio above 1.0 and a significant p-value correspond to those mutations with more decreased binding epitopes than expected. The enrichment results are in supplementary tables 2 and 3.

**Table 2:** contingency table for Fisher’s test for the enrichment of each mutation for IEDB mutated epitopes with decreased binding across 35 HLA alleles. We perform one test per mutation for each Omicron and Delta lineages.

|  | Decreased binding<br>(quotient < 0.8) | No decreased binding |
| --- | --- | --- |
| Epitopes overlapping mutation | a | b |
| Epitopes not overlapping mutation | c | d |

All the analysis and plotting code are gathered in a Jupyter notebook that is available in GitHub (https://github.com/tron-bioinformatics/omicron-analysis) together with the data and figures.

## Discussion

SARS-CoV-2 vaccines are intended to induce an immune response to the viral spike protein, including neutralizing antibodies and CD4^+^ and CD8^+^ T-cell responses. However, emergence of new variants of SARS-CoV-2 compromise the extent of protection provided by vaccines implicating the importance of T-cell responses generated by variants of concern [24]. In this study, we predicted the differential MHC binding affinity of epitopes from the emerging Omicron variant and compared it to the wide spread variant Delta across Sub-Saharan and Black South African populations. Also predicted CD8^+^ T-cell immune escape mutations based on enrichment of pMHC with decreased predicted MHC binding.

We confirmed that Omicron mutations affect a minority (19.6 %) of the epitopes reported in the IEDB database as having a positive outcome to MHC ligand or T-cell assays, as reported recently [9]. We observed that WT pMHC with strong predicted binding tend to have mutated pMHC with decreased binding in both Omicron and Delta mutations, but more markedly in the first. This observation aligns with our previous finding where such WT epitope/mutated epitope pairs show decreased predicted binding affinity [14]. However, in contrast other studies also show that SARS-CoV-2 variants of concern (VOCs) have no significant effect on CD8^+^ epitopes in COVID-19 convalescent patients [23].

Characterizing immunogenic epitopes recognized by CD8+ T-cells is challenging. It has been found that aromatic and large amino acids are enriched in immunogenic epitopes thus we hypothesize that mutations VYY143-145del, K417N and Y505H, which are replacing the aromatic Tyrosine and the large Lysine, may confer an advantage to SARS-CoV-2 [17]. Conversely, there are two mutations enriched for epitopes with non-decreased binding, T95I and D796Y, introducing the large and the aromatic amino acids Isoleucine and Tyrosine, respectively. Such mutations replacing or introducing aromatic or large amino acids are not observed in the set of Delta mutations. On the other hand, there are two, three, two and seven additional non-significant Omicron mutations introducing –instead of deleting– Phenylalanine, Tyrosine, Isoleucine and Lysine, respectively. Thus, although Omicron does contain mutations that seem to confer some level of CD8^+^ T-cell immune escape, it is unclear whether these were fixed through positive selective pressure or they are random events occurring in immunocompromised patients.

Also, the amino acid serine is negatively associated with epitope immunogenicity [17]; we reason that a mutation introducing a serine residue may confer a CD8^+^ T-cell immune escape mechanism. While no mutation in the Delta variant introduce any serine, there are two such mutations in Omicron (G446S and G496S). Nevertheless, Omicron has four additional mutations replacing a Serine. None of these is significantly enriched for decreased binding or the opposite. These spike protein mutations providing a potential CD8^+^ immune escape mechanism need to be monitored due its impact on disease severity and its effect on current global vaccination efforts. Nevertheless, Omicron does not seem to have a strong enough CD8+ immune escape and it is unclear whether through positive selective pressure such mutations may increase in future lineages of SARS-CoV-2.

## Conclusions

Our *in silico* analysis suggests that Omicron will not totally evade the immune system through a CD8^+^ immune escape mechanism. We identify some mutations that seem to be particularly relevant for CD8^+^ immune escape, but we could not confirm whether these mutations evolved through positive selective pressure. Many mutations in Omicron introduce amino acid changes associated with increased immunogenicity; this will be coherent with the hypothesis of Omicron evolution in a immuno-compromised patient.

## Acknowledgments

We thank Jonas Ibn-Salem and Franziska Lang for critical discussions. We gratefully acknowledge the authors from the originating laboratories responsible for obtaining the specimens, as well as the submitting laboratories where the sequence data were generated and shared via GISAID, NCBI Virus or the NCBI SRA, on which this research is based.

## Supplementary information

**Supplementary table 1:**
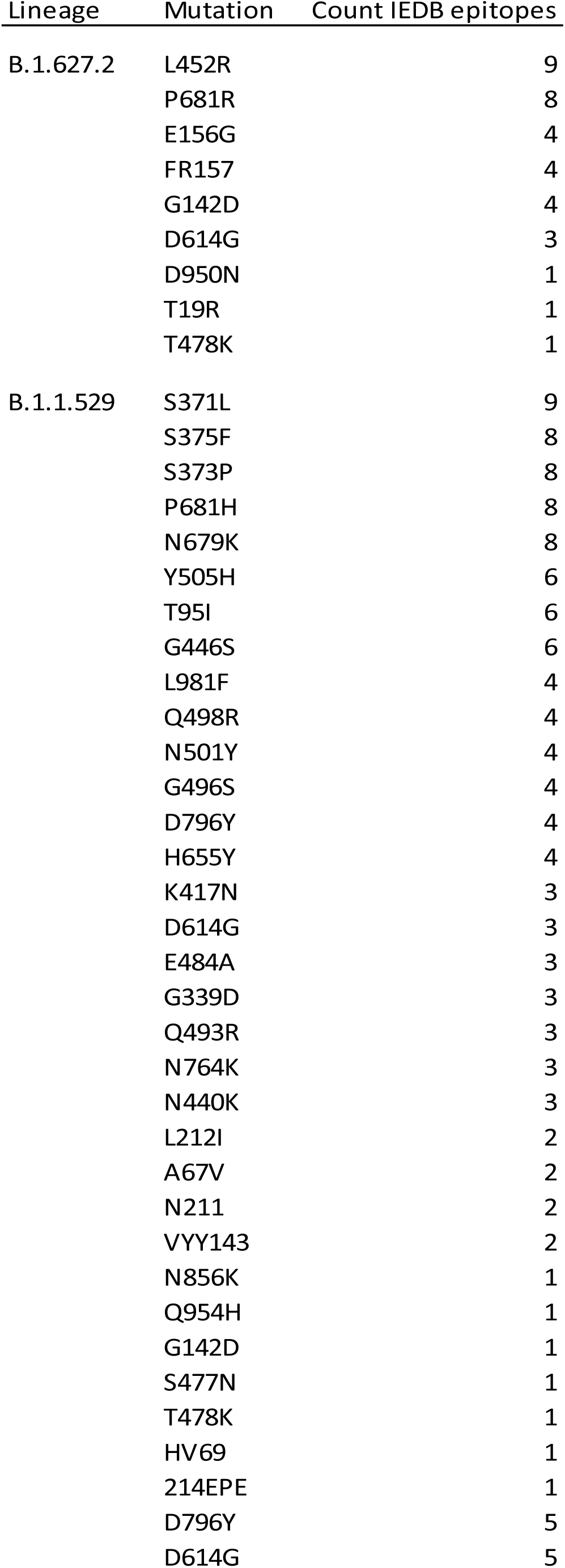
number of IEDB epitopes overlapped by each of the mutations in B.1.627.2 and B.1.1.529

**Supplementary table 2:** B.1.1.529 mutations enrichment for predicted decreased binding results

| mutation | p-value | odds ratio (OR) | a | b | c | d | adjusted p-value |
| --- | --- | --- | --- | --- | --- | --- | --- |
| N764K | 0.0003253 | 0.181001939 | 3 | 99 | 297 | 1774 | 0.000800789 |
| L981F | 0.2610271 | 0.66086573 | 11 | 102 | 289 | 1771 | 0.323098009 |
| E484A | 0.5490807 | 0.78137056 | 11 | 87 | 289 | 1786 | 0.62752081 |
| Q493R | 0.0003253 | 0.181001939 | 3 | 99 | 297 | 1774 | 0.000800789 |
| H655Y | 0.0882473 | 0.548350515 | 9 | 100 | 291 | 1773 | 0.141195682 |
| N679K | 0.2726139 | 1.226964299 | 38 | 198 | 262 | 1675 | 0.323098009 |
| P681H | 0.2726139 | 1.226964299 | 38 | 198 | 262 | 1675 | 0.323098009 |
| **T95I** | **8.84E-08** | **0.108887618** | **3** | **159** | **297** | **1714** | **4.71E-07** |
| G339D | 0.2645103 | 0.639032201 | 8 | 77 | 292 | 1796 | 0.323098009 |
| A67V | 7.97E-05 | 0 | 0 | 65 | 300 | 1808 | 0.000255131 |
| N856K | 1 | 0 | 0 | 5 | 300 | 1868 | 1 |
| **Y505H** | **9.67E-11** | **3.337348188** | **57** | **123** | **243** | **1750** | **1.03E-09** |
| **N211** | **3.74E-13** | **6.913533835** | **34** | **34** | **266** | **1839** | **5.98E-12** |
| **L212I** | **3.74E-13** | **6.913533835** | **34** | **34** | **266** | **1839** | **5.98E-12** |
| **D796Y** | **9.16E-09** | **0** | **0** | **123** | **300** | **1750** | **7.32E-08** |
| G142D | 0.236339 | 0 | 0 | 13 | 300 | 1860 | 0.323098009 |
| **VYY143** | **1.89E-07** | **5.346980976** | **21** | **26** | **279** | **1847** | **8.65E-07** |
| N440K | 0.7232497 | 0.827853881 | 8 | 60 | 292 | 1813 | 0.798068608 |
| G446S | 0.0002202 | 0.301545486 | 8 | 156 | 292 | 1717 | 0.000640469 |
| Q498R | 0.0005843 | 2.307829102 | 29 | 83 | 271 | 1790 | 0.001335575 |
| N501Y | 0.0027481 | 2.080413704 | 27 | 85 | 273 | 1788 | 0.005862622 |
| **K417N** | **2.33E-05** | **2.779320988** | **30** | **72** | **270** | **1801** | **8.29E-05** |
| **214EPE** | **5.61E-08** | **7.408244681** | **18** | **16** | **282** | **1857** | **3.59E-07** |
| **D614G** | **5.36E-06** | **3.456439394** | **25** | **48** | **275** | **1825** | **2.14E-05** |
| S371L | 0.2641312 | 0.793420652 | 32 | 245 | 268 | 1628 | 0.323098009 |
| S373P | 0.8436416 | 0.940510717 | 32 | 211 | 268 | 1662 | 0.870855846 |
| S375F | 0.8436416 | 0.940510717 | 32 | 211 | 268 | 1662 | 0.870855846 |
| G496S | 0.0139516 | 1.766182709 | 29 | 107 | 271 | 1766 | 0.027903141 |
| Q954H | 0.2483172 | 0 | 0 | 15 | 300 | 1858 | 0.323098009 |
| S477N | 0.0274157 | 0 | 0 | 30 | 300 | 1843 | 0.048738939 |
| T478K | 0.0274157 | 0 | 0 | 30 | 300 | 1843 | 0.048738939 |
| HV69 | 0.0762013 | 0.186480186 | 1 | 33 | 299 | 1840 | 0.128339003 |
| **N969K** | **3.10E-10** | **0.07** | **2** | **663** | **374** | **8158** | **1.32E-09** |
| L981F | 8.90E-01 | 0.91 | 13 | 335 | 363 | 8486 | 8.90E-01 |

**Supplementary table 3:** B.1.627.2 mutations enrichment for predicted decreased binding results

| mutation | p-value | odds ratio (OR) | a | b | c | d | adjusted p-value |
| --- | --- | --- | --- | --- | --- | --- | --- |
| L452R | 0.00011043 | 0.382415254 | 19 | 236 | 104 | 494 | 0.000496936 |
| P681R | 0.22787793 | 1.300967262 | 39 | 192 | 84 | 538 | 0.410180283 |
| G142D | 1 | 1.010377358 | 17 | 100 | 106 | 630 | 1 |
| E156G | 0.47169122 | 1.23608838 | 19 | 94 | 104 | 636 | 0.606460146 |
| FR157 | 0.47169122 | 1.23608838 | 19 | 94 | 104 | 636 | 0.606460146 |
| **D614G** | **5.99E-06** | **3.62457483** | **25** | **48** | **98** | **682** | **5.39E-05** |
| T19R | 0.80635143 | 0.784313725 | 4 | 30 | 119 | 700 | 0.907145364 |
| D950N | 0.1465259 | 0 | 0 | 15 | 123 | 715 | 0.329683279 |
| T478K | 0.01490101 | 0 | 0 | 30 | 123 | 700 | 0.044703024 |

**Supplementary figure 1:**
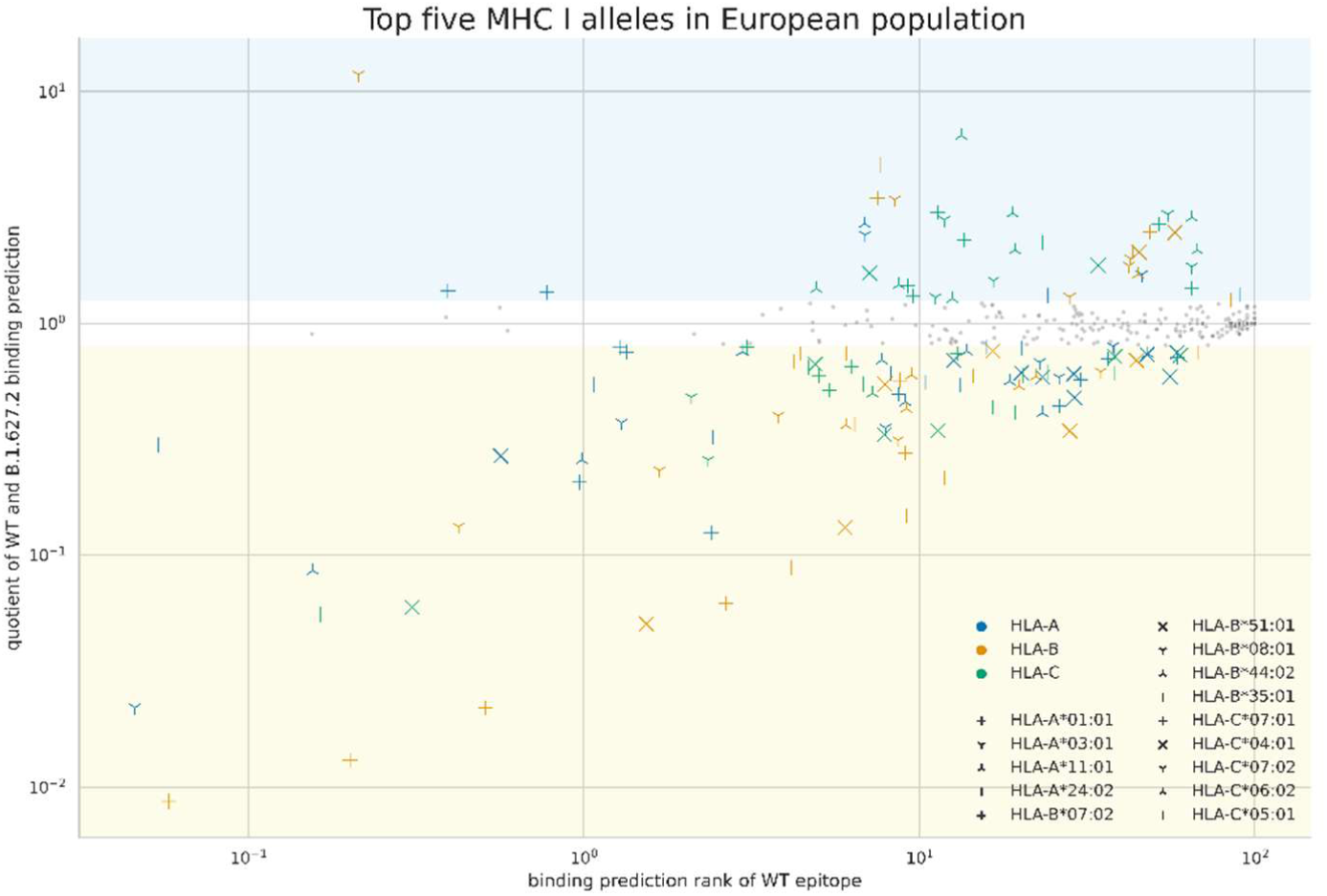
differential MHC binding of the IEDB epitope-homologous epitope pairs in B.1.627.2 on top five HLA alleles on the European population.

**Supplementary figure 2:**
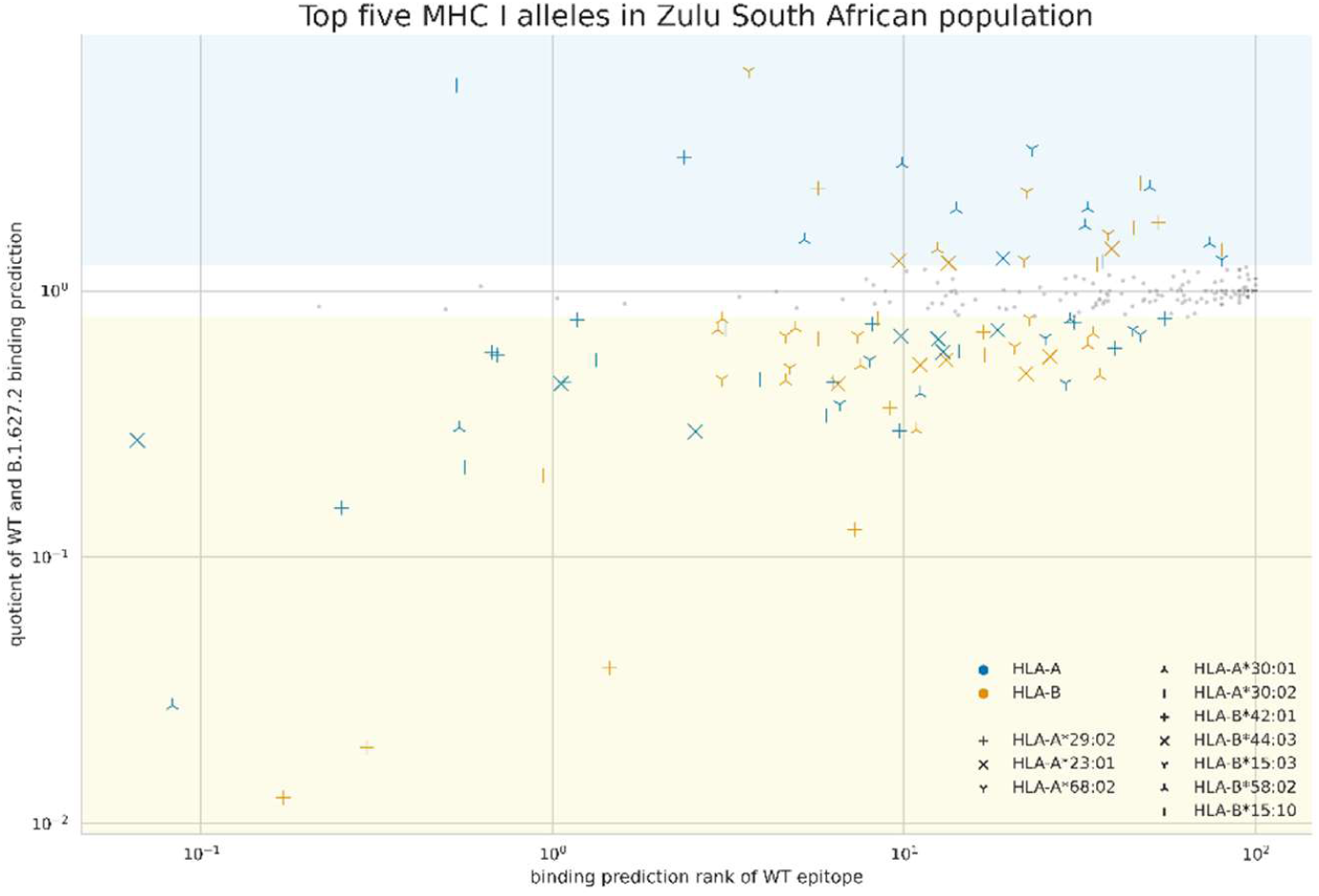
differential MHC binding of the IEDB epitope-homologous epitope pairs in B.1.627.2 on top five HLA alleles on the Zulu South African population (only HLA-A and HLA-B available).

**Supplementary figure 3:**
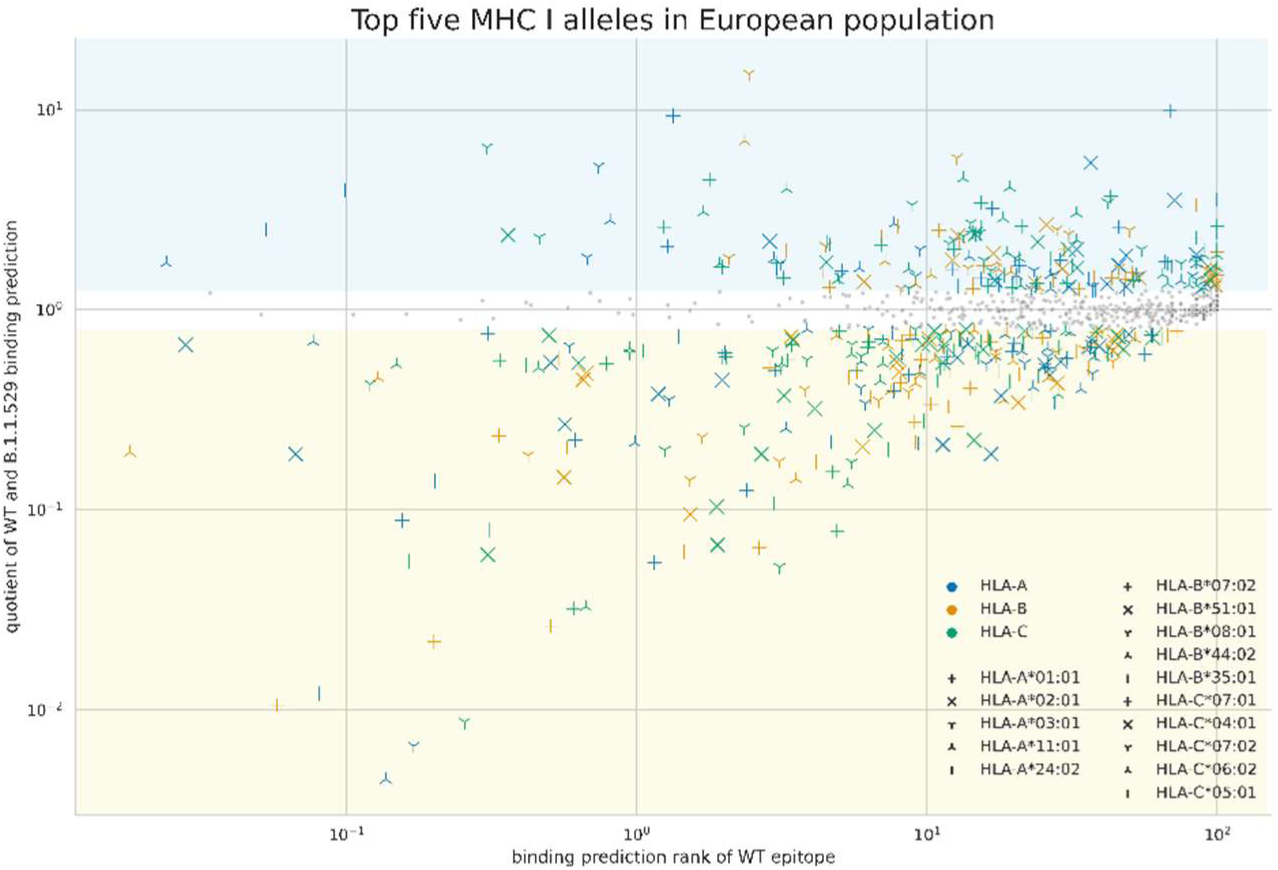
differential MHC binding of the IEDB epitope-homologous epitope pairs in B.1.1.529 on top five HLA alleles on the European population.

**Supplementary figure 4:**
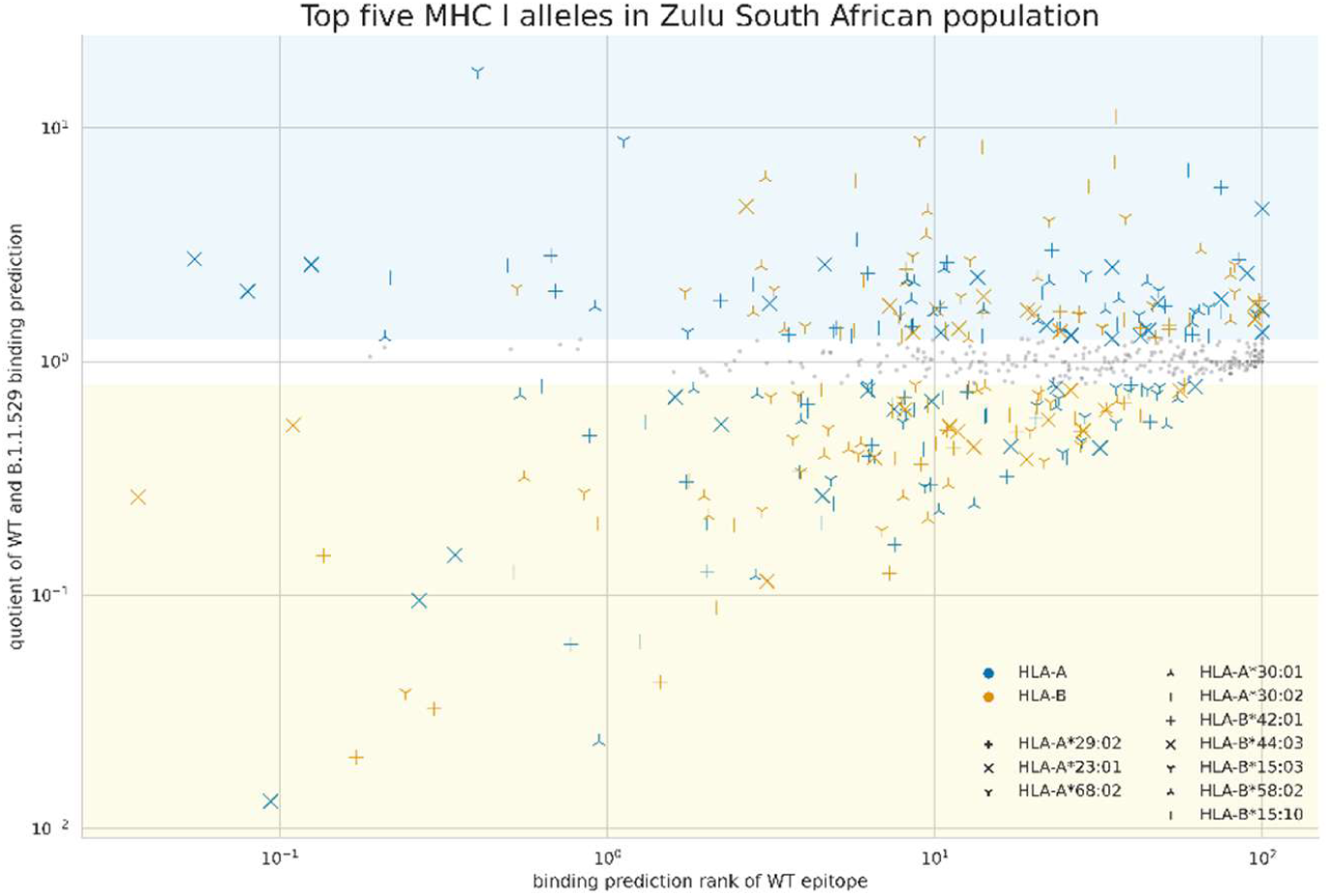
differential MHC binding of the IEDB epitope-homologous epitope pairs in B.1.1.529 on top five HLA alleles on the Zulu South African population.

**Supplementary figure 5:**
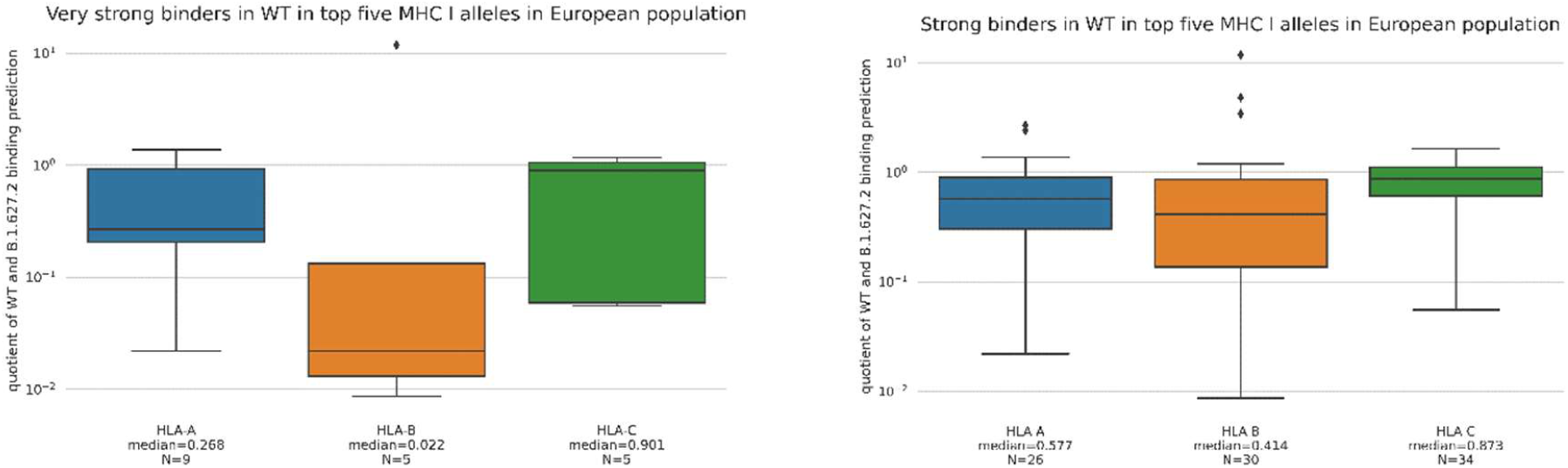
distribution of quotient of predicted MHC binding in the WT and B.1.627.2 for strong and very strong binders in the top five HLA alleles in the European population.

**Supplementary figure 6:**
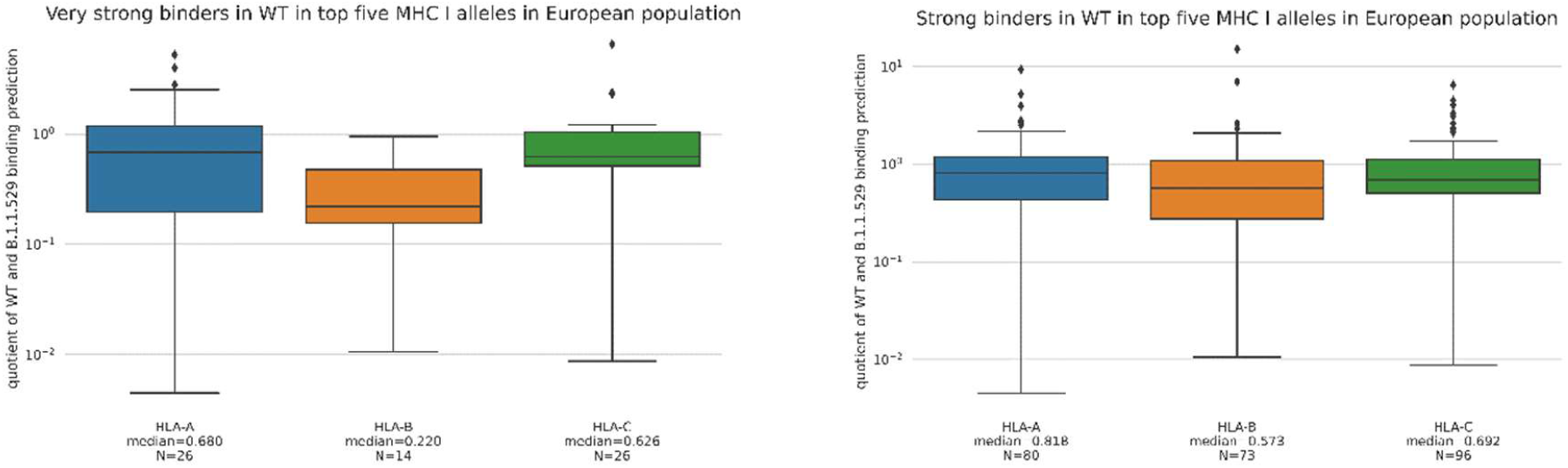
distribution of quotient of predicted MHC binding in the WT and B.1.1.529 for strong and very strong binders in the top five HLA alleles in the European population.

